# ENTRAIN: integrating trajectory inference and gene regulatory networks with spatial data to co-localize the receptor-ligand interactions that specify cell fate

**DOI:** 10.1101/2023.07.09.548284

**Authors:** Wunna Kyaw, Ryan C. Chai, Weng Hua Khoo, Leonard D. Goldstein, Peter I. Croucher, John M. Murray, Tri Giang Phan

## Abstract

Cell fate is commonly studied by profiling the gene expression of single cells to infer developmental trajectories based on expression similarity, RNA velocity, or statistical mechanical properties. However, current approaches do not recover microenvironmental signals from the cellular niche that drive a differentiation trajectory. We resolve this with environment-aware trajectory inference (ENTRAIN), a computational method that integrates trajectory inference methods with ligand-receptor pair gene regulatory networks to identify extracellular signals and evaluate their relative contribution towards a differentiation trajectory. The output from ENTRAIN can be superimposed on spatial data to co-localize cells and molecules in space and time to map cell fate potentials to cell-cell interactions. We validate and benchmark our approach on single-cell bone marrow and spatially resolved embryonic neurogenesis datasets to identify known and novel environmental drivers of cellular differentiation. ENTRAIN is available as a public package at https://github.com/theimagelab/entrain and can be used on both single-cell and spatially resolved datasets.

## Main text

In multicellular organisms, cells in different organs and tissues adopt different states of cellular differentiation to allow them to perform specialized tasks. The precise coordination of cellular differentiation and function requires not only the existence of multiple distinct cellular fates but also the ability of the cells to communicate and regulate each other to maintain homeostasis and avoid disease^1^. The development of single-cell technologies such as single-cell RNA sequencing (scRNA-seq) has revolutionized our ability to deconvolute the myriad of heterogenous cellular transcriptional states that comprise multicellular life, even in seemingly homogenous cell lineages such as natural killer (NK) cells^2^. Interestingly, scRNA-seq has suggested that cells exist in a continuum of transcriptional states, whereas the traditional assignment of cell identity by the expression of cell lineage markers, such as by flow cytometry, have viewed cell fates as discrete, non-overlapping entities^3^. Thus, the cell state is the transcriptional output of the gene regulatory networks and may represent transient intermediate steps in the differentiation of the cell towards its developmental destination, or cell fate^4, 5^. Accordingly, it may also be possible to predict the future cell fate from the current cell state and the dynamic expression of critical master regulator genes.

Trajectory inference computes the pattern of change in gene expression for cells in a given dataset and arranges them in pseudo-chronological order along a developmental pathway (pseudotime) based on the similarity between their changing gene expression profiles^6, 7^. There are currently more than 70 published trajectory inference methods, with many more in development^6^. This reflects both the popularity of pseudotime for lineage tracing and also the limitations of the technique, which are dependent on the underlying assumptions, many of which are project and cell-type specific^8^. RNA velocity is an alternative approach that uses the relative abundance of unspliced to spliced mRNA transcripts to predict future cell states, instead of inferring them from global similarity in the transcriptomic profiles between cells^9, 10^. However, the modelling of RNA kinetics also makes several assumptions, such as a common rate of splicing across different genes and the sampling of multiple intermediate cell states in addition to the mature steady-state^11^. The RNA velocity analysis of peripheral blood mononuclear cells (PBMCs), which contain mature blood cells without the immature bone marrow precursor cells, is a good example of the potential for this approach to generate spurious cell lineage relationships^11, 12^. Thus, there are fundamental limits to the fidelity of dynamic inferences that can be made from single cell snapshots^13^. The cross-validation of cell state transitions and lineage relationships by additional orthogonal methods has therefore been strongly recommended^11, 12^.

The development of tools for ligand-receptor (LR) network analysis of single cell data has made it possible to decipher the cell-cell communications that may also drive cell state transitions and determine cell fate^1^. First used to infer cellular interactions at the feto-maternal interface in the human placenta^14^, LR analysis has become increasingly popular with its ability to infer interactions between cells in a given dataset, even in the absence of spatial information^15^. Broadly, tools for LR analysis can be generalized into two categories: 1. ‘LR-only’ tools that rely solely on ligand-receptor gene expression, and 2. ‘LR + Intracellular’ tools that incorporate intracellular regulons. ‘LR-only’ tools, such as CellPhoneDB^16, 17^, predict cell-cell interactions by considering the expression of ligand and receptor genes as a proxy for secreted and membrane protein abundance. Tools from the ‘LR + Intracellular’ category are motivated by the possibility that a scarcely expressed LR pair may also unexpectedly regulate a considerable array of downstream genes, which would be overlooked by ‘LR-only’ tools that only consider gene expression levels. To this end, these tools exploit the large body of biological prior knowledge about gene regulatory networks and intracellular signalling pathways to prioritize LR pairs based on their downstream influence on gene regulation. As a result, methods belonging to the ‘LR + Intracellular’ category achieve markedly different results from methods in the ‘LR-only’ category. Thus, LR analysis has potential to complement trajectory inference and RNA velocity by providing corroborating evidence for gene regulatory programmes responsible for cell state transitions. However, only two tools belong to the second category, NicheNet^18^ and CellCall ^19^, and no tools to date incorporate trajectory or velocity information with LR analysis.

The introduction of spatially resolved transcriptomics has demonstrated the important role of physical location within a tissue. Specifically, different stages of differentiation within a population often correlate with microanatomical location in the tissue^20^. Similarly, LR interactions are limited by surface contact between interacting cells, or through diffusivity for secreted ligands^21^. This suggests that the spatial information of a cell, which is typically lost in traditional scRNA-seq workflows, can improve the evaluation of LR pairs that influence the differentiation trajectories of a cell. Therefore, there is a need for computational methods that incorporate spatially resolved data to better understand the environmental drivers of differentiating populations.

Here, we have integrated the information provided by trajectory inference and RNA velocity with LR analysis to develop ENTRAIN, an environment-aware trajectory inference computational tool that can be used to predict the extracellular drivers of cell state transitions. ENTRAIN consists of three modules, ENTRAIN-Pseudotime, ENTRAIN-Velocity, and ENTRAIN-Spatial, which can be applied on the outputs of pseudotime-based methods, RNA velocity or paired single-cell and spatially resolved data, respectively. In turn, ENTRAIN can be applied to a wide range of datasets containing differentiating cells as well as the cell’s interacting microenvironment, including spatial datasets. The ENTRAIN package is available to download at https://github.com/theimagelab/entrain.

## METHODS

### Materials and Methods

#### Assumptions and Overview

ENTRAIN operates based on certain assumptions about the biological system of interest:

1. Environmental control over a differentiating cell population, if present, is facilitated through LR interactions.
2. The environmental influence on differentiation is operating on a time scale resolvable by either pseudotime-based or RNA velocity methods.
3. The environmental regulation occurs via known regulatory pathways that are documented in gene regulatory network databases, and that the degree of regulation in this database can be quantified as the edge weight (*w*) between a given ligand (*l*) and a given gene *g* ∈ *G*, where *G* denotes the set of all genes in the genome.

The fundamental operating principle of ENTRAIN is that, if a specific ligand *l*, is influencing the expression of a specific gene *g* in a differentiating population, this influence can be observed as a meaningful contribution of the ligand-gene regulatory network towards predicting the observed changes in the expression of *g*. In other words, if the edge weight *w* between *l* and *g*. which represents the strength of the regulatory interaction, positively correlates with the observed gene expression changes in the trajectory (or velocity), then this suggests that the ligand is actively driving the observed differentiation for that gene in the observed dataset.

First, we construct differentiation trajectories either by using manifold-based trajectory inference tools^7^ or RNA velocity estimation with scVelo^10^. We then identify trajectory informative (‘TRAINing’) genes that either correlate their expression with pseudotime (for manifold-based trajectories) or exhibit high velocity likelihoods (for RNA velocity-based methods). In parallel, we identify LR pairs using NicheNet^18^ and extract regulatory interactions between identified LR pairs and downstream target genes in the regulon. We then fit a random forest regression model using TRAINing gene covariances (for pseudotime) or velocity probabilities (for scVelo) as the ‘response’ variable and NicheNet predicted regulatory interactions as the ‘explanatory’ variable. This model estimates the proportion of trajectory dynamics (as measured by pseudotime covariance or velocity likelihood) that can be predicted by the regulatory interactions downstream of a LR pair. Ligands are scored based on their contributions to the model.

### Trajectory construction with Monocle

Consider cells as *n*, vectors in ℝ^|*G*|^, where |*G*| is the number of genes measured by the scRNA-seq experiment and *n* is the number of cells. Typically, a differentiation process will take the form of an ordered sequence of cells in this high dimensional space, beginning at a root cell (or node), traversing along a series of intermediate cells with progressive changes in gene expression before ending at a terminal cell. In this ordered sequence, called pseudotime, cells that are highly similar in gene expression space will be adjacent in pseudotime. Assuming sufficient sampling of intermediate cell stages, this approach successfully identifies differentiation trajectories but cannot determine whether a trajectory is driven by its environment or is under cell-intrinsic control, motivating the use of ENTRAIN to identify environmental influences. ENTRAIN implements pseudotime analysis by using the Monocle3^22^ workflow, which applies the SimplePPT^23^ tree algorithm to cells in reduced dimension space to calculate cell pseudotimes (*τ*_i_,…, *τ*_n_).

### Selection of TRAINing genes

Because trajectory pseudotime *τ* is derived from underlying gene expression profiles, we hypothesized that a trajectory can sufficiently be described by several trajectory informative TRAINing genes: driver genes whose expression levels exhibit strong linear relationships with pseudotime, and presumably have a greater influence on pseudotime calculation and graph learning. Biologically, we assume that genes with strong linear relationships with pseudotime are highly significant in differentiation processes. Specifically, consider a single trajectory branch ***B***, consisting of an *n*, cells by |*G*| genes expression matrix:

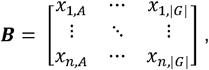

Where *n* is the number of cells in ***B***, *A* denotes a gene, and *x*_l,*A*_ denotes the expression of gene *A* in cell 1. Each cell (1,…,*n*) has a corresponding pseudotime (*τ*_I_,…, *τ*_n_,). We aim to identify influential TRAINing genes by using gene-pseudotime covariance as a metric for evaluating gene significance in a differentiation trajectory:

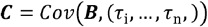

In each branch, genes are ranked by covariance and the lowest ranked genes (default: bottom 5%) are removed from the workflow to prevent these from confounding further analysis. The remaining genes are classified as TRAINing genes for that trajectory.

We note that TRAINing genes are distinct from commonly used ‘differentially expressed genes’ in two ways: 1. TRAINing genes are not dependent on cell type annotations, and 2. TRAINing genes may not necessarily exhibit large absolute changes in expression as one traverses a cell lineage but strongly co-vary with pseudotime. It is this covariance, rather than absolute expression, that is used to define TRAINing genes.

While covariance is the default metric, ENTRAIN can alternatively be configured to use correlation coefficients.

### Extracting regulatory information from NicheNet

Expression dynamics during differentiation are likely to be a manifestation of cell-intrinsic and cell-extrinsic regulatory programmes. To demarcate these two factors, the algorithm’s second step unites prior knowledge of ligand-receptor pairs and their corresponding intracellular regulatory interactions to determine potential ligands driving the observed TRAINing gene expression dynamics.

Under the assumption that the microenvironmental niche has a quantifiable contribution to gene expression dynamics in differentiation, we require a database that predicts which target genes are subject to regulation by ligand-receptor pairs. ENTRAIN extracts this information from NicheNet ^24^, which unites traditional ligand-receptor signalling to downstream transcriptional regulation. We first identified active LR pairs amongst the trajectory cells (‘receivers’) and the remaining cells in the dataset (‘senders’), using NicheNet as prior knowledge of possible ligand-receptor interactions. With the assumption that high LR expression levels do not necessarily correlate to significance in driving differentiation trajectories, we determined LR pairs for further analysis if they fulfilled two criteria: 1. They are expressed by a sufficient proportion of cells in the dataset (default >0 counts in at least 10% of cells). 2. The corresponding receptors are expressed by a sufficient proportion of differentiating cells (default >0 counts in at least 10% of differentiating cells). Of the ligands that meet the criteria, we extracted their respective downstream target regulation scores from the NicheNet database. These are vectors representing the ability of a given ligand to regulate every human gene. Thus, each ligand is associated with a vector of length *g*, where *g* is the number of human genes in the database, and each element of the vector is a number (a “regulatory potential”) representing the strength of the regulatory relationship between the ligand and a given gene.

### Calculation of top environmental drivers of a trajectory

Next, we assumed that some subset of the active ligands will constitute the extracellular signals influencing a trajectory. We speculated that the regulation between ligands and the trajectory could be contained in existing databases of regulatory networks interactions.

To detect this, we used a supervised random forest model to fit NicheNet regulatory potentials (explanatory variable) to TRAINing gene covariances (response variable) 25. Here, we consider the NicheNet matrix as an *L* by |*G*| matrix ***L***, where *L* is the number of actively signalling ligands, and the covariances are represented by a |*G*| dimensional column vector ***C***. Random forest attempts to fit ***L***, to ***C***, used with hyperparameters *n_trees* = 500, *n_features at each split* = number of ligands (features) divided by 3.

In principle, some columns of ***L*** (which represent the predicted change in gene expression as a result of the ligand-receptor pairing), will possess greater similarity to ***C*** than others if the ligand is responsible for the observed covariance in ***C***. This similarity is represented as variable importance, calculated by removing one column at a time from the matrix and calculating the loss in Gini index that results from the removal. Thus, variable importance represents the significance of a ligand in predicting observed gene expression covariance.

To assess the environmental dependence of whole trajectory branches, we used % Variance Explained (%V.E.). This metric measures how well the random forest predicts the variance in ***C***. More formally,

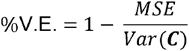

Random forest was chosen as the primary algorithm for feature scoring owing to several advantages suited for our context. Firstly, it caters to non-linear interactions between features, such as those that might be found in regulatory interactions between ligands and their downstream target genes. Secondly, built-in methods for feature selection and scoring, based on sequential removal of features, accommodates our primary goal of scoring ligands rather than predicting gene expression. Thirdly, while a known drawback of random forests is the difficulty of interpretability, this is offset by our existing prior knowledge of gene regulatory networks that provides the insight into downstream targets. Lastly, considering the relatively low numbers of ligands and receptor genes relative to the rest of the genome, the computational complexity of random forests compared to other feature selection algorithms becomes less concerning. Moreover, our fitting is performed on the level of trajectory branches or velocity clusters, rather than individual cells, further mitigating concerns of computational complexity.

### Calculation of cell-wise influences

Differentiating cells exhibit changes in receptor expression and regulatory wiring as they progress along a developmental process. Because of this, we hypothesized that certain stages of a developmental process will be more influenced by environmental signalling than other stages. We thus wished to produce a more granular, cell-wise measure of ligand influence that encapsulates this behaviour. To do this we calculated pseudotime-expression covariances along a rolling window of cells along pseudotime, restricted to separate branches (**Supplementary Algorithm 1**). We used a default window size *w* and step size *s* of 10% and 2% of the cells in the trajectory branch, respectively. This ‘local covariance’ quantifies a gene’s expression dynamics within a rolling window of differentiating cells. To this end, we fit a second round of random forest models to each rolling window, such that every branch is now subject to an additional 50 ‘local’ model fits corresponding to 50 rolling windows along the branch. The number of local model fits is dependent on the values of *s* and *w*; 50 rolling windows is the behaviour when *s* and *w* are assigned default values. We used regulatory potentials from the top 5 ligands as the predictor variable (a |*G*| *x* 5 matrix with default parameters) and the local covariances as the response variable. For step sizes greater than 1, we linearly interpolate %V.E._*i*_ values for cells which are skipped.

Resultant %V.E. values denote the confidence of the NicheNet fit at each of the 50 windows. Genes possessing high covariance with pseudotime are assumed to be important for trajectory determination, and we are interested in the subset of those that are under environmental control. Some of these high-covariance genes will not be under extracellular control and consequently exhibit a low %V.E. value when fitted to NicheNet. On the other hand, high covariance genes that are also under extracellular control will exhibit both high covariances and a confident fit (increased %V.E.) to NicheNet. As a result, these window %V.E. values can be interpreted as the degree of environmental dependence across different stages of the trajectory. Ultimately, every trajectory branch is subject to one ‘branch-wide’ model fit that determines the top few ligands of interest, and 50 ‘local’ model fits that assess where their regulatory effects are most noticeable. Cells with cell-intrinsic drivers would be expected to exhibit low, negative, or widely varying %V.E. values as the model cannot accurately fit environmental regulators to the observed expression dynamics in that window, while the opposite is true for highly environmentally dependent windows. We note that the term ‘cell-wise’ is slightly misleading, as the observed expression dynamics are deduced from the covariances of many neighbouring cells in a rolling window of observations rather than a single cell.

### Finding ligands responsible for RNA velocity dynamics

RNA velocity is a dynamical approach that calculates the time-derivative of RNA concentration for single cells, allowing for short-term predictions of cell fate in differentiating populations. Because these dynamics are often dependent on environmental signals, we predicted that ENTRAIN could be employed to determine driver ligands responsible for observed RNA velocity vectors. Biologically, these represent ligands that may be responsible for short time scale dynamics that may not be resolvable using the pseudotime-based approach described previously. For full details of the velocity estimation, see ref. ^26^.

In most datasets, a small minority of genes are responsible for the majority of observed velocity variance^27^, necessitating a way to prioritize velocity genes by their significance. The ENTRAIN-Velocity module uses scVelo to recover fit likelihoods, a measure of velocity significance ^26^, from which to infer ligand activity (**Supplementary Figure S1**).

We first clustered the RNA velocity matrix into *c*, groups representing major axes of variance in RNA velocity vectors, by repurposing the Leiden algorithm in scanpy^28^. We then calculated the fit likelihoods for velocity genes, by applying the scVelo recover_dynamics^26^ function to each velocity cluster. For each velocity cluster *c*_*i*_, this process generates a vector ***ℓ***_***i***_ of length |*G*_*i*_|, where|*G*_*i*_ | is the number of genes with calculated fit likelihoods per cluster *c*_*i*_. Note that the genes with calculated fit likelihoods are usually a subset of all genes because not all genes possess confident velocities. These genes (row names) constitute our TRAINing genes for this module, and the fit likelihoods (values) represent the response variable for subsequent model fit described below.

To elucidate environmental influence driving the velocities, we fit the NicheNet ligand-target matrix to all genes with calculated likelihoods using a random forest regression model ^25^ with hyperparameters *n_trees* = 500, *n_features at each split* = number of ligands (features) divided by 3. As before, we consider the NicheNet matrix as an *L* by |*G*_*c*_| matrix ***L***, and the velocity likelihoods for a given cluster *c*_*i*_ are represented by a |*G*_*c*_| dimensional column vector ***ℓ***_***C***_. Random forest attempts to fit ***L*** to ***ℓ***_***i***_ for all clusters *c* (**Supplementary Algorithm 2**), under the assumption that if a ligand is truly responsible for some component of the observed velocities in a cluster, the corresponding column in ***L*** will be more similar to the velocity likelihood vector compared to less significant ligands. Similarly to the pseudotime-based approach, we extracted mean decrease in Gini index and %V.E. scores to evaluate ligand significance.

### Finding ligands responsible for RNA velocity dynamics in spatially resolved datasets

The third module of ENTRAIN, called ENTRAIN-Spatial, is designed for datasets with paired scRNA-seq and Visium data. This module first calculates and clusters velocities on the scRNA-seq matrix object, as in ENTRAIN-Velocity. This is followed by transferring velocity cluster labels to the Visium dataset using the package tangram-sc^29^. Next, within each velocity cluster, the ENTRAIN-Spatial subsets the Visium dataset to include only those spots matching the velocity cluster label or the spots in direct adjacency.

Subsequently, we select genes that are included in NicheNet’s ligand-receptor network to inform later analysis of ligand-receptor pairings. In contrast to the previous ENTRAIN-Velocity module, these genes are restricted to those that are situated in the immediate spatial vicinity of differentiating cells.

Subsequent ligand-receptor pairing, random forest fitting, and scoring were performed identically as in the ENTRAIN-Velocity module.

## RESULTS

ENTRAIN explicitly incorporates output from established trajectory tools to inform a random forest feature selection model for ligand scoring (**Fig. 1A)**. As a proof-of-concept, we validated ENTRAIN on a scRNA-seq dataset profiling the bone marrow microenvironment (BME) and its resident mesenchymal and haematopoietic lineages in mice (**Fig. 1B**). We evaluated the contribution of each gene towards the trajectory dynamics by calculating pseudotime using Monocle3 (**Fig. 1C**). We extracted the gene expression for cells along the pre-B trajectory (**Fig. 1D**) and derived the pseudotime-expression covariance for every gene. We assessed the biological relevance of this metric by ranking the genes by covariance and interrogating the top covarying TRAINing genes. This revealed known lineage marker genes including *Vpreb1, Igll1*, and *Vpreb3* for pre-B cells. In parallel, we examined the microenvironmental interactions by selecting receptor and ligand genes. We then queried the NicheNet ligand-target regulatory potential database to obtain regulatory interactions between active ligands and their corresponding regulons. ENTRAIN was then performed on the developing B cell lineages using this database as input. We calculated the model’s V.E., a measure of the proportion of TRAINing gene covariance that can be attributed to extracellular signals. The percentage of V.E. by the 71 identified active ligands was 2.6%. To identify more granular behaviour, we conducted ENTRAIN in a cell-wise manner by analysing environmental dependence in a series of 100 rolling windows along trajectory pseudotime for every branch. This analysis revealed that the previous environmental dependence was restricted to small pockets of HSCs (**Fig. 1E**), indicating that the local ligand influence was restricted to a subpopulation of progenitor cells that appeared relatively early in lineage commitment. ENTRAIN output shortlisted signalling ligands that are known to be involved in B cell development (*Vcam1/Lama2-Itgb, Il7-Il7r*; *Tnfsf13b-Tnfrsf13b; Il15/Il2-Il2rg*) and ligands with conserved roles during cellular differentiation (*Dll1-Notch1/2; Dkk2-Lrp6*; *Jag1-Notch1/2*) (**Fig. 1F**). The regulatory potential was dominated by a small subset of functionally relevant target genes, particularly *Ebf1, Myl4* and *Cd79a*. Interrogating the source of these ligands revealed that while some of the top-ranked ligands were expressed primarily by a singular cell type (**Fig. 1F**, coloured lines), others were expressed among heterogenous cell types (grey lines). ENTRAIN also identified a novel extracellular signal that was not previously known to be involved in B cell development (*Ptdss1-Scarb1/Jmjd6*) (**Fig. 1F**).

**FIGURE 1:**
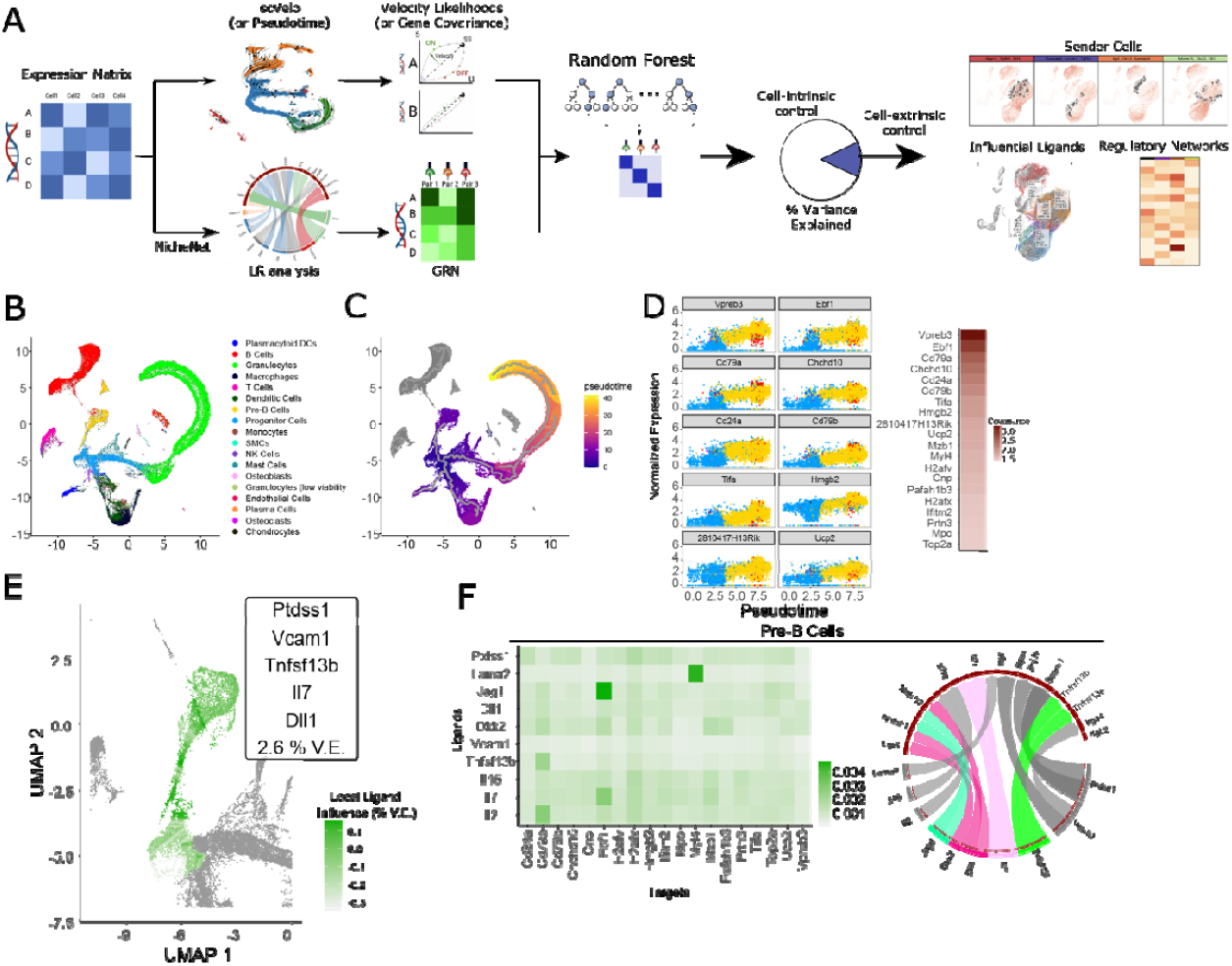
ENTRAIN-Pseudotime analysis of pre-B cell development. (A) ENTRAIN workflow. (B) UMAP representation of 133,942 cells in mouse bone marrow environment. (C) Monocle3 trajectory overlayed on the UMAP. (D) High trajectory covariance (TRAINing) genes for the trajectory between haematopoietic progenitors and pre-B cells. (E) ENTRAIN ligand results overlayed on the B cell lineage trajectory. Cells coloured by local ligand influence. V.E: Variance Explained (F) Ligand-target gene regulatory networks (left) and circos plot (right) representing regulatory links between top ranked ligands and their downstream targets. Colour represents identity of major cell type expressing that ligand. Ligands expressed by more than one cell type are coloured grey.

To demonstrate the versatility of ENTRAIN we developed the ENTRAIN-Velocity module to recover environmental signals responsible for the RNA velocity vector and applied it to a murine embryonic neurogenesis dataset^30^ (**Figure 2A)**. The velocity matrix was recovered using scVelo and clustered with the Leiden algorithm^31^ to deconvolute velocity variance into major groups. The vectors formed 10 velocity clusters (VC0-9), which roughly corresponded to major cell types and transitions (**Figure 2B**). We analysed and ranked the joint likelihoods of the velocities in each cluster to identify the TRAINing genes for this dataset: the most rapidly up-or down-regulated genes during neurogenesis (**Fig. 2C**). We then applied ENTRAIN to each velocity cluster to identify driver ligands responsible for the observed velocities (**Fig. 2D**). The analysis predicted positive V.E. scores for 5 out of 10 clusters (VC0-VC3 and VC7) corresponding to velocities exhibited by fibroblastic, radial glial, neuroblast/neuronal, and neural tube cell clusters (**Fig. 2E**). In these clusters, the environmental influence was attributed to ligands in the *Notch* pathway (*Tgfb2, Bmp2, Ntf3* and *Bdnf*) and *Wnt* signaling pathway (*Sema3b, Psap* and *Pdgfb*) known to be involved in embryonic neurogenesis. More generally, we considered ligands ranked among the top 5 in each positive cluster and showed that 21 out of 25 ligands were known to be involved in embryonic neurogenesis (**Supplementary Table S1**), with the exceptions being the extracellular matrix proteins *Npnt*/*Adam15* and *Serpinc1*. Interrogation of the NicheNet ligand-target network revealed interactions between *Tgfb2*-*Ina*/*Mapt*/S*tmn2/Igfbpl1, Bdnf-Bcl11b, and Jag1-Ebf1* as major components of environment-driven neuronal differentiation (vcluster1 and vcluster7), as well as *Jag1-Sdc2* as the largest environmental driver in mesenchymal development (vcluster2) (**Fig. 2F**). Fibroblasts and neuroblasts were the major cell types responsible for producing the highest 3 ranked ligands (**Fig. 2F**).

**FIGURE 2:**
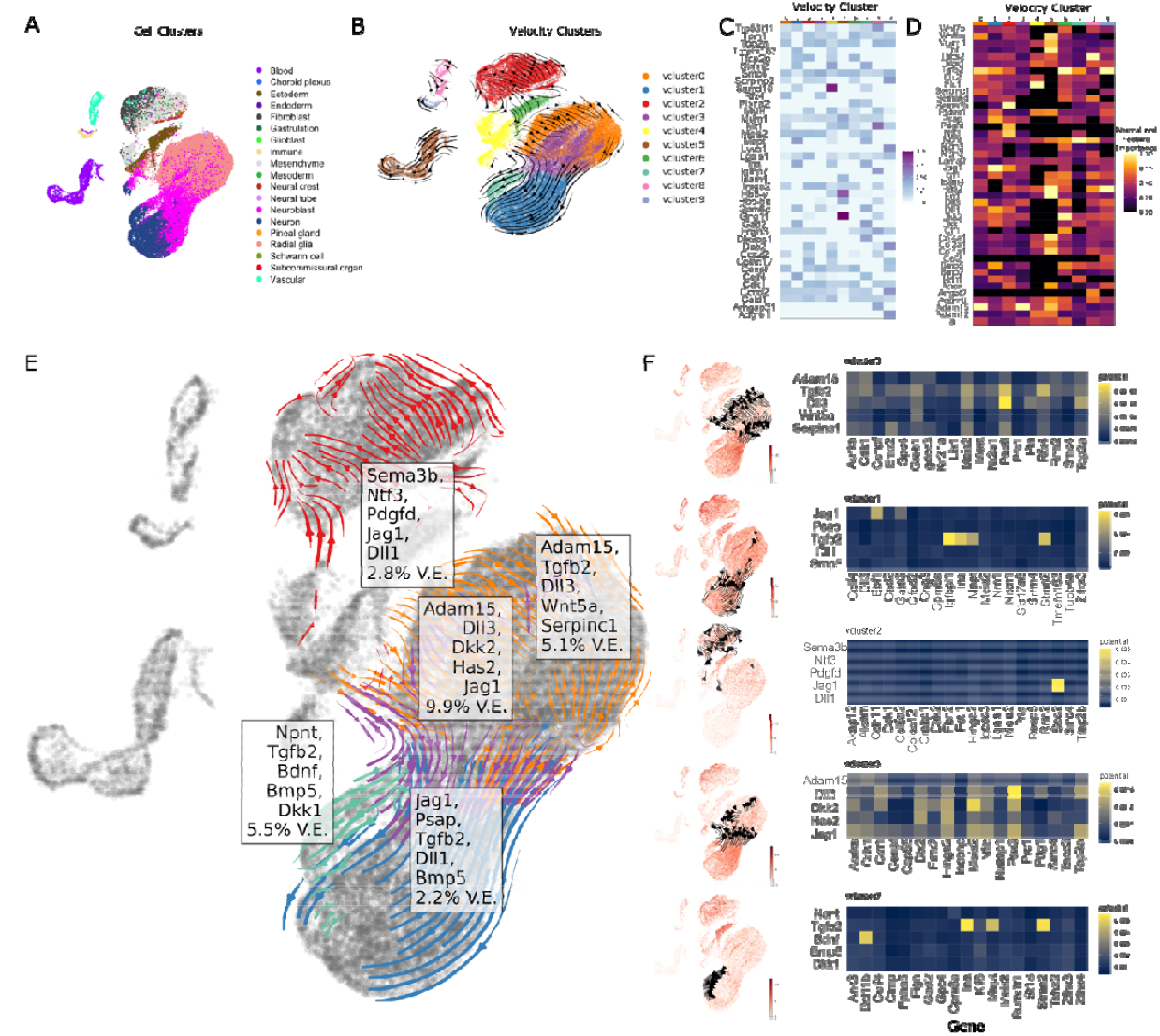
ENTRAIN-Velocity analysis of neuronal development. (A) UMAP representation of mouse embryonic neurogenesis dataset at E10.5. (B) Velocity vectors overlayed on UMAP representation, cells coloured by velocity cluster membership. (C) Heatmap of high likelihood velocity genes in each velocity cluster. (D) Heatmap of ligands predicted by ENTRAIN to influence velocities in each velocity cluster. (E) Velocity vectors, V.E. scores and top 5 ligands predicted by ENTRAIN for 5 out of 10 clusters (VC0-3, VC7) overlayed on the UMAP embedding. (F) Sender expression and predicted gene targets for the top 5 ligands in each velocity cluster. Left: UMAP embedding coloured by mean expression of the top 5 ligands predicted for the velocity cluster. Right: Heatmap showing NicheNet regulatory linkages between the top 5 ligands (y-axis) and their downstream target genes (x-axis).e

Emerging spatial transcriptomics technologies have recently shown success in delineating the role of cell-cell communication in various cellular contexts^21, 32^. Building upon this, we developed the ENTRAIN-Spatial module to decode cell-cell communication signals driving RNA velocities, while concurrently considering their spatial environment. This module operates by accepting a paired dataset of spatial transcriptomics data and single-cell data. Its output comprises those ligand-receptor pairs that are both spatially co-localized and have a quantifiable influence on the observed RNA velocities.

We applied ENTRAIN-Spatial to a paired dataset consisting of both 10x Chromium single-cell and 10x Visium data, which was obtained from Ratz et al.^33^ (**Fig. 3A** and **Fig. 3B**). We recovered the RNA velocities from the 10x Chromium data using scVelo^26^ and subsequently clustered the velocities into 8 major clusters (**Fig. 3C**). By utilizing Tangram^29^, we transferred the velocity cluster labels to their spatial positions.

**FIGURE 3:**
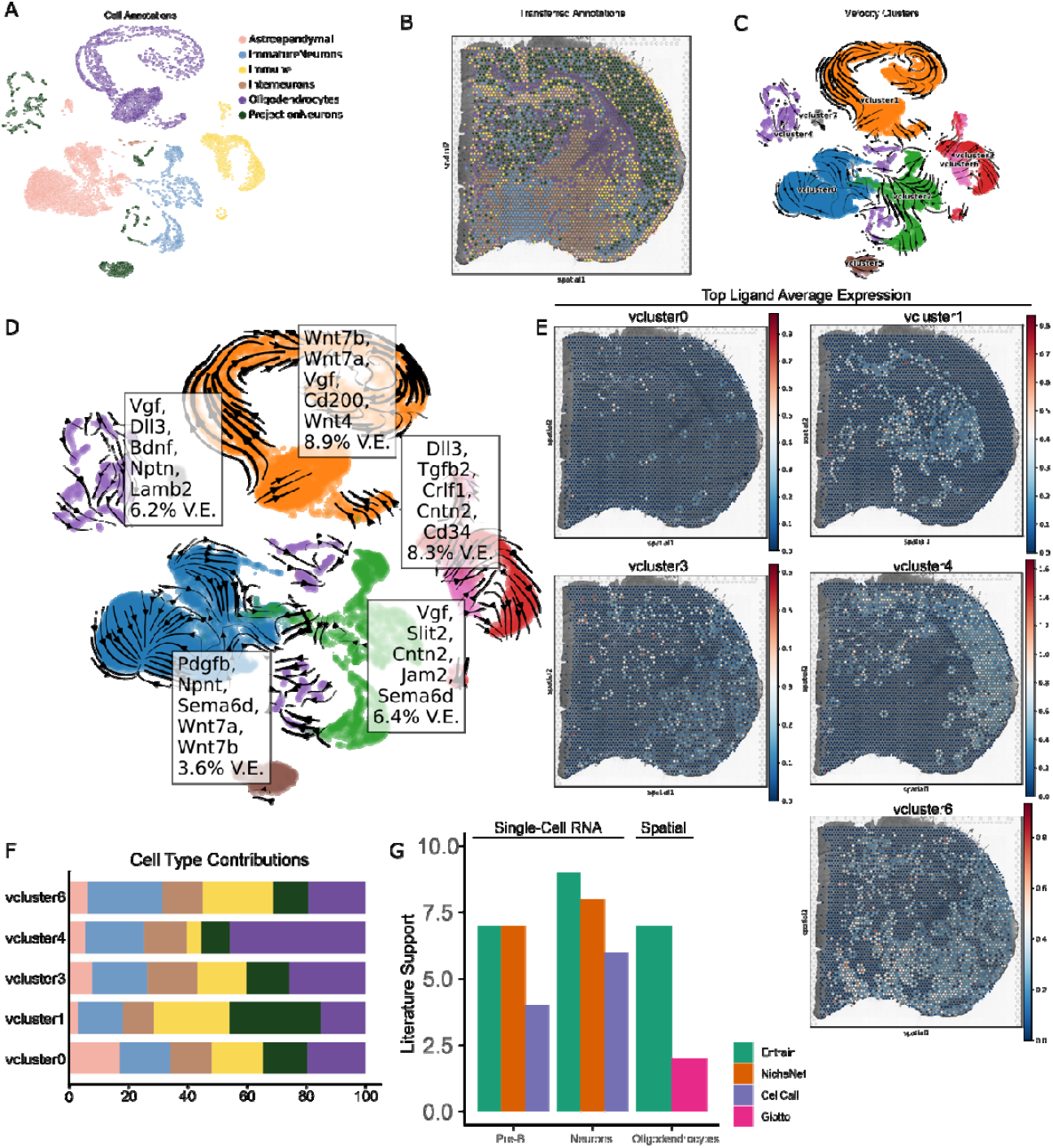
ENTRAIN-Spatial analysis of neuronal development at spatial resolution. (A) UMAP plot of pre-annotated Ratz et al. dataset (B) Tangram transferred labels overlayed on spatial scatter plot. (C) UMAP plot of velocity cluster labels. (D) Top 5 ligands predicted by ENTRAIN for positive V.E. clusters (Velocity Clusters 0, 1, 3, 4 and 6) overlayed on velocity plot. (E) Spatial scatter plot representing average expression of top 5 ligands associated with each velocity cluster. (F) Stacked bar plot showing the proportion of cell types expressing the top 5 ligands for each velocity cluster, weighted by variance explained. (G) Bar plot showing number of ligands with literature support for their role in pre-B cell and neuronal development.

We then used ENTRAIN-Spatial to evaluate ligands located in close spatial proximity to spots associated with a specific velocity label. The scoring was performed based on each ligand’s potential to instigate the observed RNA velocities. ENTRAIN-Spatial results indicated that five out of the eight major velocity clusters (vcluster0, 1, 3, 4 and 6) exhibited a detectable level of environmental influence, as quantified by the percentage of variance explained (% V.E.) (**Fig. 3D**). Notably, the velocity cluster corresponding to immature and mature oligodendrocytes (vcluster3) demonstrated the highest proportion of variance explained. This cluster corroborated ligands that are well-documented to be implicated in oligodendrocyte maturation, including the *Wnt*-family and *Vgf*. Notably, as opposed to ENTRAIN-Velocity, these ligands are restricted to those expressed in any spot adjacent to a spot associated with a velocity cluster.

To interpret spatial patterns in driver ligand expression, ENTRAIN-Spatial facilitates the visualization of specific spots expressing the highest-ranking ligands (**Fig. 3E**) as well as the relative contributions of spatially adjacent cell types towards driving the observed velocities (**Fig. 3F**).

To corroborate our findings, we benchmarked the performance of ENTRAIN to similar methods NicheNet^18^ and CellCall^19^ for single-cell RNA results, and Giotto^34^ for spatial transcriptomics results, concentrating specifically on the top 10 ligands from each method, as well as the highest velocity confidence clusters (**Supplementary Figure S2**), to maintain. Despite the observed discrepancy between all these methodologies (**Supplementary Fig. S3**), ENTRAIN demonstrated the highest rate of literature support across the top ranked ligands when analyzing the pre-B cell, neuroblast, and oligodendrocyte lineages (**Fig. 3G**).

These results indicate that ENTRAIN accurately recovers extracellular regulators that are not resolved by DEG-based methods.

## DISCUSSION

ENTRAIN uses an orthogonal approach that has several advantages over other methods that are highly dependent on the accurate identification of DEGs, which in turn depend on correct and reproducible cell type clusters, labels and pair-wise comparisons. As a result, these methods cannot consider intra-cluster expression dynamics that may arise as a cell differentiates along a trajectory. In comparison, ENTRAIN can be executed on any arbitrary number of cell states linked by a trajectory or RNA velocity vectors. In turn, ENTRAIN can analyse sparse populations that are not amenable to DEG-based methods.

ENTRAIN exhibits several limitations. Firstly, ENTRAIN-Pseudotime is dependent on the quality of the topology that is learnt by the trajectory inference algorithm^35^. To mitigate this, the ENTRAIN-Pseudotime module allows flexible input from any trajectory method provided that each input cell is assigned a pseudotime value and a trajectory branch in the Seurat object metadata. In addition, ENTRAIN allows interactive selection of trajectory nodes for flexible analysis on a user-defined branch. Secondly, ENTRAIN-Velocity is similarly subject to the same limitations as RNA velocity. Namely, the potential for inferring spurious velocity vectors when it is applied to populations with multiple kinetic regimes or datasets containing mature cell types missing intermediate cell states^12^. Thirdly, the NicheNet database does not discriminate between up- or down-regulated targets, which may result in ENTRAIN detecting both inhibitors and activators of a differentiation pathway. Lastly, ENTRAIN requires whole-transcriptome based technologies to ensure accurate capture of all ligand and receptor genes. Therefore, hybridization-based technologies which detect a limited panel of genes may not be suitable.

In conclusion, we present ENTRAIN, the first tool to date that integrates trajectory and cell-cell communication methods to identify driving ligands influencing cell differentiation. Validating ENTRAIN on existing single-cell pre-B Cell, neuronal, and spatially resolved brain datasets demonstrates that ENTRAIN recovers cell-extrinsic determinants of differentiation. Comparative analysis suggests that ENTRAIN outperforms other cell-cell communication methods in deciphering intercellular signals governing differentiation, possibly owing to the leveraging of trajectory and velocity data rather than traditional differential expression. Future work may consist of extension towards capturing epigenetic contributions from methylation or chromatin accessibility data^36, 37^

## Acknowledgments

We thank Joakim Lundeberg, Joseph Powell for discussions. This project is supported by the UNSW Cellular Genomics Futures Institute and the Australian National Health and Medical Research Council (NHMRC) Ideas Grant APPID2003662. T.G.P. and P.I.C. and this work are supported by Mrs. Janice Gibb and the Ernest Heine Family Foundation. P.I.C. (APPID2009010), R.B. (APPID2008356), and T.G.P. (APPID1155678) are supported by Investigator Grants and Fellowships from the NHMRC. L.D.G. is supported by the Kinghorn Foundation. W.K. is supported by an Australian Government Research Training Scholarship.

