## Supplemental Information for "ENTRAIN: integrating trajectory inference and gene regulatory networks with spatial data to co-localize the receptor-ligand interactions that specify cell fate"

**SUPPLEMENTARY METHODS**

**Supplementary** **Algorithm 1:**

**Inputs**

Trajectory branches: **b**, a collection of ($N$ x $|G|$ ) cell by gene matrices b_i_, one for each branch (pre-dimension reduction). Each cell is assigned a pseudotime $\tau_{i}$

NicheNet Ligand Target Database**:** $\boldsymbol{L}$ ($L$ x $|G|$) matrix

$s$: step size, an integer denoting the interval (in # of cells) between successive rolling windows.

$w$: window width, an integer denoting the size (in # of cells) of the rolling window.

**For each** branch b_i_ in **b**

Sort cells in b*_i_* by pseudotime $\tau_{i}$

**for** $\text{(}m\text{ ≤ }N\text{)}$,

Select a window of cells $(p, \ldots, m, \ldots, q)$, where $p = m-\frac{w}{2}, q = m+\frac{w}{2}$, and $m$ denotes the window centre.

Calculate covariances $\boldsymbol{C}$ between pseudotimes $\left( \tau_{p},\ldots, \tau_{m} , \ldots, \tau_{q} \right)$ and gene expression in the window. Specifically,

$\boldsymbol{C=}Cov\left( \left[ \begin{matrix} x_{p,a} & \cdots& x_{p,|G|} \\ \vdots& \ddots& \vdots\\ x_{q,a} & \cdots& x_{q,|G|} \end{matrix} \right], \left( \tau_{p},\ldots,\tau_{m} , \ldots, \tau_{q} \right) \right)$, where $x_{m,a}$denotes the gene expression of gene $a$ in cell $m$

Fit $\boldsymbol{L}$ to $\boldsymbol{C}$ with randomForest

Store ${\text{\%}\text{V.E.}}_{i}$ value from randomForest for the window centre cell $m$ in a vector $\text{R}$

$m\text{ = }m\text{ + }s$

**if** ($s>1$)

Linearly interpolate values of $\text{R}$ for all cells that were not assigned as a window centre. For example, If $s$ = 5, every 5^th^ cell will be denoted a window centre, meaning that 4 out of every 5 cells in the dataset will be missing an ${\text{\%}\text{V.E.}}_{i}$ value. These 4 cells will have their values interpolated.

**Supplementary** **Algorithm 2:**

**Inputs**

$\mathbf{V}$**:** Velocity matrix, a ($N$ x $|G_{v}|$) cell by gene matrix denoting RNA velocities.

$\boldsymbol{L}$ **:** NicheNet Ligand Target Database ($L$ x $|G_{v}|$ matrix, where $L$ is the number of active ligands present in the cell population

**P:** Leiden clustering resolution parameter (0 $\leq$ $P$ $\leq$ 1)

1. Cluster $\mathbf{V}$ using Leiden algorithm with resolution $P$, resulting in $c$ clusters and a corresponding number of matrices $\mathbf{V}_{\boldsymbol{i}}$ of shape $N_{i}$ x ${|G}_{v}|$, where $N_{i}$ is the number of cells in a given cluster $c_{i}$

**For each** $c_{i}$ in $c$**:**

1. Calculate gene likelihoods using scvelo.tl.recover_dynamics() on $\mathbf{V}_{\boldsymbol{i}}$**,** the corresponding velocity matrix for cells in cluster $c_{i}$. This generates a likelihoods vector $\mathcal{l}_{\boldsymbol{i}}$ of length ${|G}_{i}|$, where ${|G}_{i}|\leq{|G}_{v}|$, containing likelihoods calculated for genes denoted as velocity genes.
2. Define $\boldsymbol{L}_{\boldsymbol{i}}\boldsymbol{=}\boldsymbol{L}_{\boldsymbol{j}}\boldsymbol{| row(}\boldsymbol{L}_{\boldsymbol{j}}\boldsymbol{) \in}\mathcal{l}_{\boldsymbol{i}}\boldsymbol{,} \forall\boldsymbol{j}$**.** In other words, $\boldsymbol{L}_{\boldsymbol{i}}$ consists of rows from $\boldsymbol{L}$ that correspond to genes with calculated fit likelihoods.
3. Fit $\boldsymbol{L}_{\boldsymbol{i}}$ to $\mathcal{l}_{\boldsymbol{i}}$ with randomForest.
4. Append ${\%\text{V.E.}}_{i}$ (% velocity variance explained by ligands for cluster $c_{i}$) and $i_{l}$ (vector of length $L$, denoting variable importance) to vectors $\boldsymbol{R}$ and $I_{l}$.

**return:** $\boldsymbol{R}$, $I_{l}$

**Benchmarking and Validation**

To select the clusters to benchmark against, we conducted a velocity confidence analysis using scVelo to identify the most reliable velocities for comparison. In both the Manno and Ratz datasets, velocity clusters corresponding to neuroblasts/neurons and oligodendrocytes, respectively, exhibited the highest confidence among clusters with positive variance explained (**Supplementary Figure S2**). As a result, our benchmarking efforts focused on these specific velocity clusters for comparison against both the literature and other methods.

**Benchmarking against NicheNet**

NicheNet was applied to the BME/EN datasets using default parameters. Sender cells were set as ‘undefined’ to determine LR pairs in an agnostic manner. “Condition_oi” (condition of interest) and “condition_reference” were defined as Pre-B Cells and Progenitor cells (for Pre-B cell differentiation), or “Neuron” and “Neuroblast” (for Neuroblast differentiation). NicheNet results were then ranked by Pearson correlation to determine the ranking order.

**Benchmarking against CellCall**

CellCall was applied to the BME/EN datasets using default parameters provided in documentation. CellCall results were then filtered to ligand-receptor interactions involving the cell type of interest (i.e., Pre-B cells or neuroblasts). Ligands were then ranked by scores to determine the ranking order.

**Literature Curation**

Because NicheNet and ENTRAIN return a scored list of all active ligands, without defining an ‘active ligand’, we compared the top 10 ligands from each method for literature curation.

For each ligand, a literature search was performed following these specifications:

1) The ligand has been mentioned to be involved in extracellular signalling
2) The ligand has been reported to be involved in differentiation or development of the cell type of interest: Pre-B cell development, or neurogenesis, or oligodendrocyte differentiation.
3) Ligands reported to be involved regulation in the cell type of interest but not in differentiation (e.g., lymphoblastic lymphoma, glioblastoma multiforme), or involved in non-developmental differentiation (e.g., post-germinal centre B cell differentiation) were excluded.

**EXPERIMENTAL DETAILS**

**Mice**

6- to 8-week-old immunocompetent C57BL6/J mice. Animal experiments were approved by the Garvan Institute of Medical Research Animal Ethics Committee (ARA16/01 and ARA19/09).

**Isolation of endosteal and marrow cells**

To obtain bone and bone marrow stroma cells for scRNA-seq, mice were sacrificed via CO2 asphyxia. Femurs were harvested and soft tissue was removed. The femurs were then separated into diaphysis and metaphysis (epiphysis was removed). Marrow cells were collected by flushing the diaphysis with PBS. Endosteal cells were isolated from marrow-depleted diaphyseal and metaphyseal bone fractions by gently crushing and cutting bones and digested in 2 mg/ml of collagenase A and 2.5mg/ml of trypsin for 30 mins at 37ºC. After digestion, bone fractions were vortexed for 10s and the supernatant containing digested cells was filtered through a 100μm filter into collection tubes containing 10% FCS. Marrow cells and endosteal cells were then centrifuged at 400x g for 5 mins and resuspended in 200uL and were then stained in PBS supplemented with 2% FCS for FACS sorting.

**FACS enrichment of endosteal and marrow cells**

Cells were stained for Ter119-PE at 4ºC for 30mins and rinsed with PBS supplemented with 2% FCS. Dead cells and debris were excluded by FSC, SSC and DAPI (ThermoFisher Scientific). Cells that were viable (DAPI-negative) and negative for erythroid marker (Ter119) were sorted into PBS supplemented with 2% FCS.

**Single cell RNA-seq**

Single cells were encapsulated into emulsion droplets using the 10x Chromium (10x Genomics). scRNA-seq libraries were constructed using Chromium Single Cell 30 v2 Reagent Kit according to the manufacturer’s protocol. Briefly, FACS sorted sample volume was decreased and cells were examined under a microscope and counted with a cell counter (Thermo Fisher Scientific). Cells were then loaded in each channel with a target output of 10,000 cells. Reverse transcription and library preparation were performed on C1000 Touch Thermal cycler with 96-Deep Well Reaction Module (Bio-Rad). Amplified cDNA and final libraries were evaluated on a Agilent BioAnalyzer using a High Sensitivity DNA Kit (Agilent Technologies). Individual libraries were diluted to 4nM and pooled for sequencing. Pools were sequenced with 75 cycle run kits (26bp Read1, 8bp Index1 and 55bp Read2) on the Novaseq Sequencing System (Illumina) to 80%-90% saturation level.

**Pre-processing of 10x scRNA-seq data**

Raw sequencing data were processed using the CellRanger pipeline (10x Genomics). Count matrices were loaded into R and further processed using Seurat^1^. We removed all cells with fewer than 300 distinct genes observed or cells with more than 10% of unique molecular identifiers stemming from mitochondrial genes.

**Dimensionality reduction**

Dimensionality reduction was performed using gene expression data for the top 3000 variable genes. The variable genes were selected based on dispersion of binned variance to mean expression ratios using FindVariableGenes function of Seurat package^1^. Next, principal component analysis (PCA) was performed and the first 40 principal components were included for subsequent clustering and UMAP analysis based on manual inspection of a principal component variance plot (‘PC elbow plot’).

**Clustering and sub-clustering**

Graph-based clustering of the PCA reduced data with the Louvain Method was performed^2^ after computing a shared nearest neighbor graph^3^. The clusters were visualized on a 2D map produced with Uniform Manifold Approximation and Projection (UMAP)^4^. For sub-clustering, we applied the same procedure of finding variable genes, dimensionality reduction, and clustering to the restricted set of data (usually restricted to cell clusters of the same lineage).

**Differential expression of gene signatures**

The marker genes for each cluster were identified using the FindAllMarkers function and ROC-based test statistics of Seurat.

**Filtering out doublets**

It is to be expected that a fraction of data should consist of cell doublets (and to an even lesser extent of higher order multiplets) due to co-encapsulation into droplets and/or pairs of cells that were not dissociated in sample preparation. Therefore, clusters of cells expressing markers of different lineages were removed from further analysis.

**SUPPLEMENTARY FIGURES**

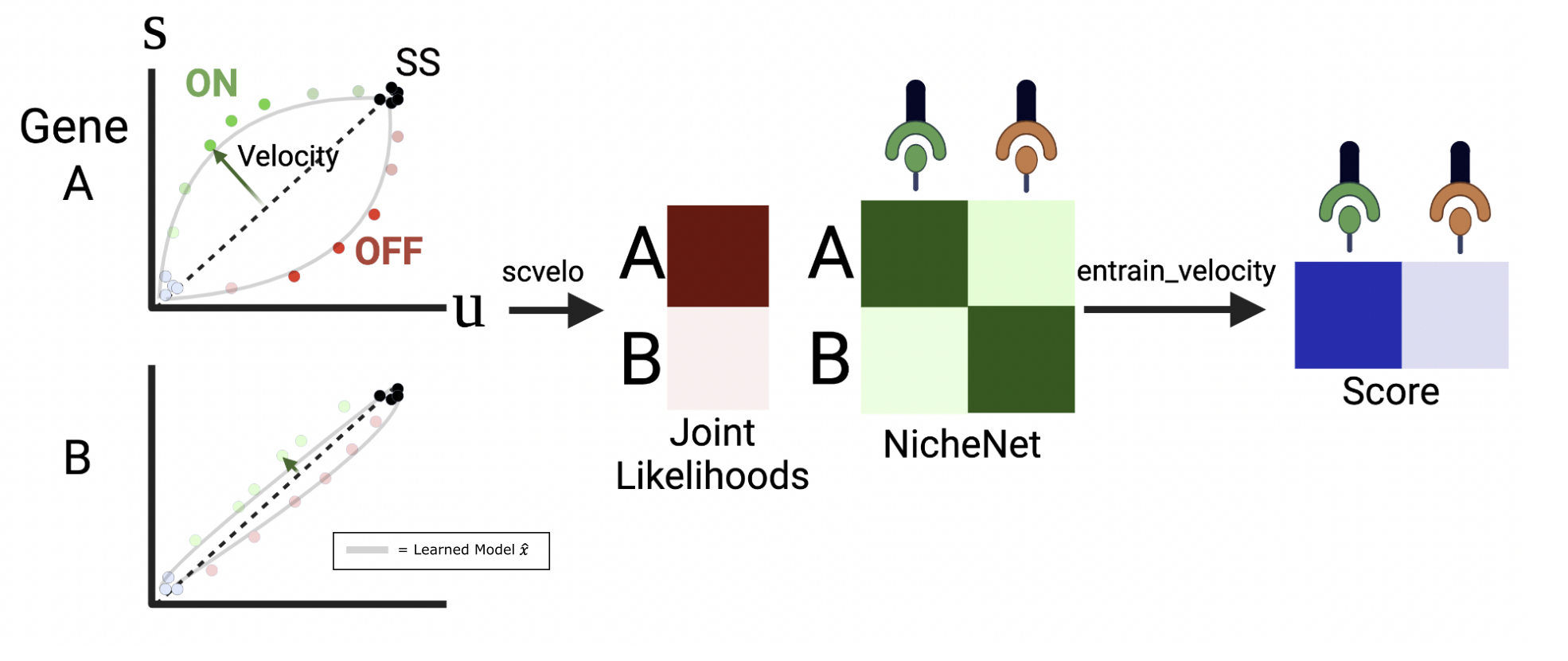

**Supplementary Figure S1:** For a given gene, scVelo fits a phase trajectory model $\hat{x}$ to the un-spliced and spliced counts. Left, scatter plot: Cells are assigned to states on the phase diagram based on their distance to the phase model, with likelihoods of the assignment denoted by color opacity (black = steady-state ON, green = ON/upregulation, red = OFF/downregulation, light blue = steady-state OFF). Middle: Likelihoods across all cells are pooled into a joint likelihood for each gene (red squares). Right: ENTRAIN fits the NicheNet matrix to the likelihoods vector according to similarity between the velocity likelihoods and predicted ligand-gene relationships (green squares), resulting in a variable importance score (blue) for each ligand that denotes the significance of a ligand in driving the observed velocities.

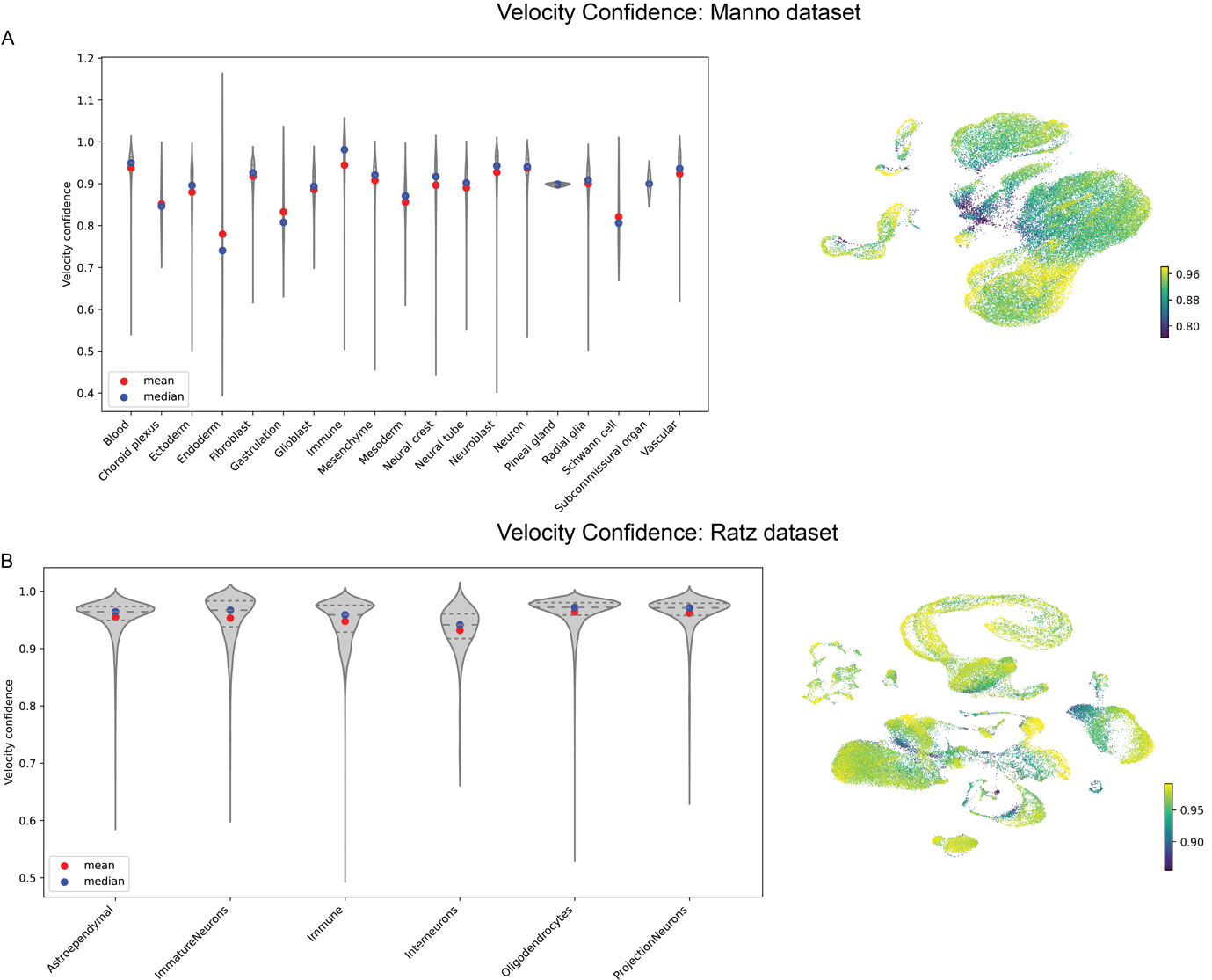

**Supplementary Figure S2:** Velocity confidence analysis of Manno (A) and Ratz (B) datasets per cell type cluster (left) and at single-cell (right) resolution.

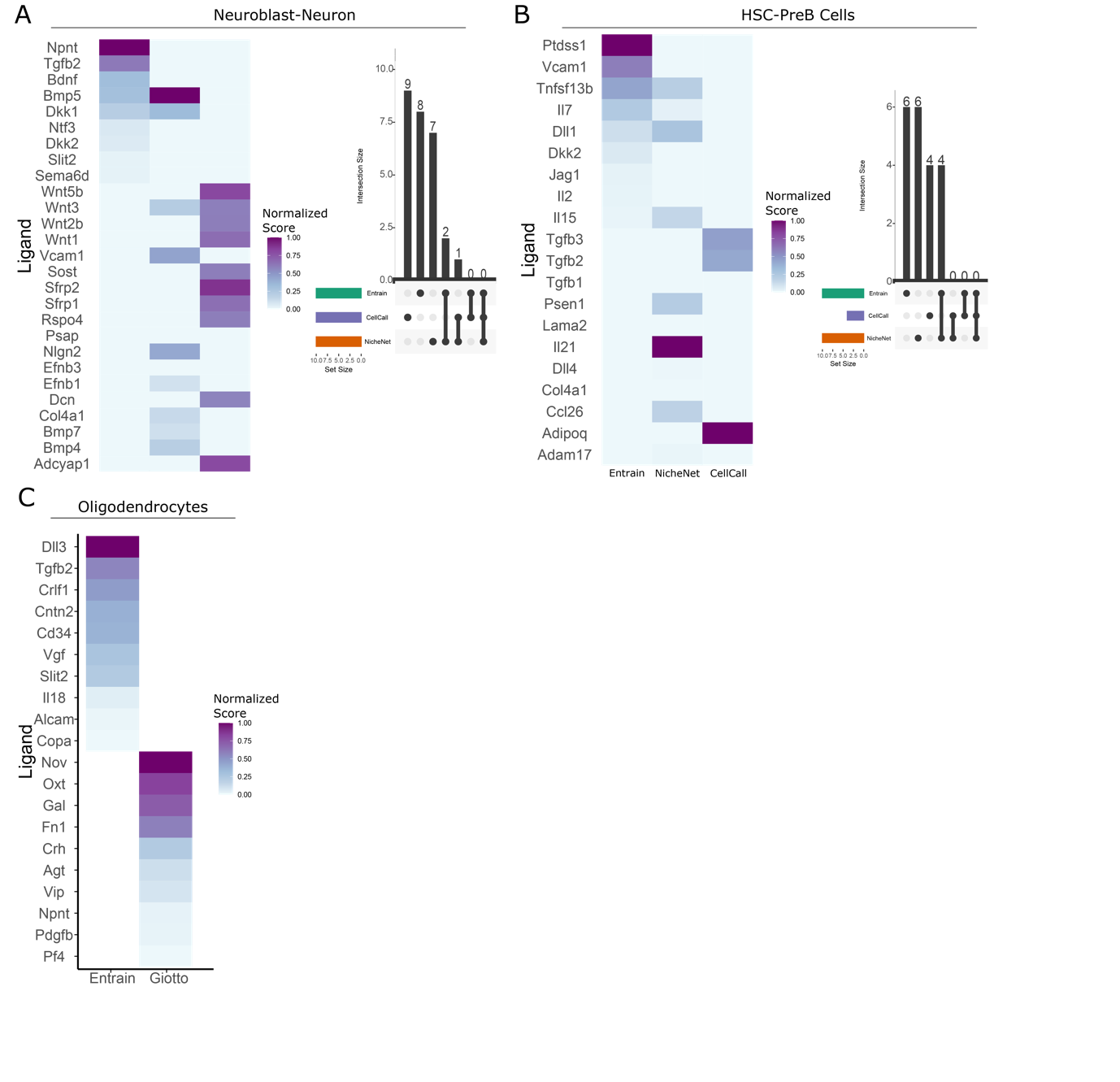
**Supplementary Figure S3:**
(A) Left: Heatmap showing normalized scores of the top 10 ligands found by ENTRAIN, NicheNet and CellCall performed on the neuroblast-neuron subpopulation in EN. Right: UpsetR plot showing overlap between the ligands predicted by each method.

(B) As in (A) but for Progenitor-Pre-B Cell subpopulation in BME.

(C) Heatmap showing normalized scores of the top 10 ligands found by ENTRAIN and Giotto performed on the oligodendrocyte subpopulation of the Ratz et al. dataset
